# Establishment of the family Zarkiaceae (Oscillatoriales, Cyanobacteria) and description of the new marine genera *Zarkia* (Zarkiaceae, Oscillatoriales) and *Romeriopsis* (Leptolyngbyaceae, Synechococcales), from northern Portugal

**DOI:** 10.1101/2021.04.09.439031

**Authors:** Guilherme S. Hentschke, Ângela Pinheiro, Vitor Ramos, Aldo Barreiro, M. Sofia Costa, Sébastien Brule, Vitor M. Vasconcelos, Pedro N. Leão

**Affiliations:** Interdisciplinary Centre of Marine and Environmental Research (CIIMAR/CIMAR), University of Porto, Av. General Norton de Matos, 4450-208, Matosinhos, Portugal; Centro de Investigação de Montanha (CIMO), Instituto Politécnico de Bragança, Campus de Santa Apolónia, 5300-253 Bragança, Portugal; Faculty of Sciences, University of Porto, Rua do Campo Alegre, 4169-007 Porto, Portugal

**Keywords:** subtidal, intertidal, phylogeny, taxonomy

## Abstract

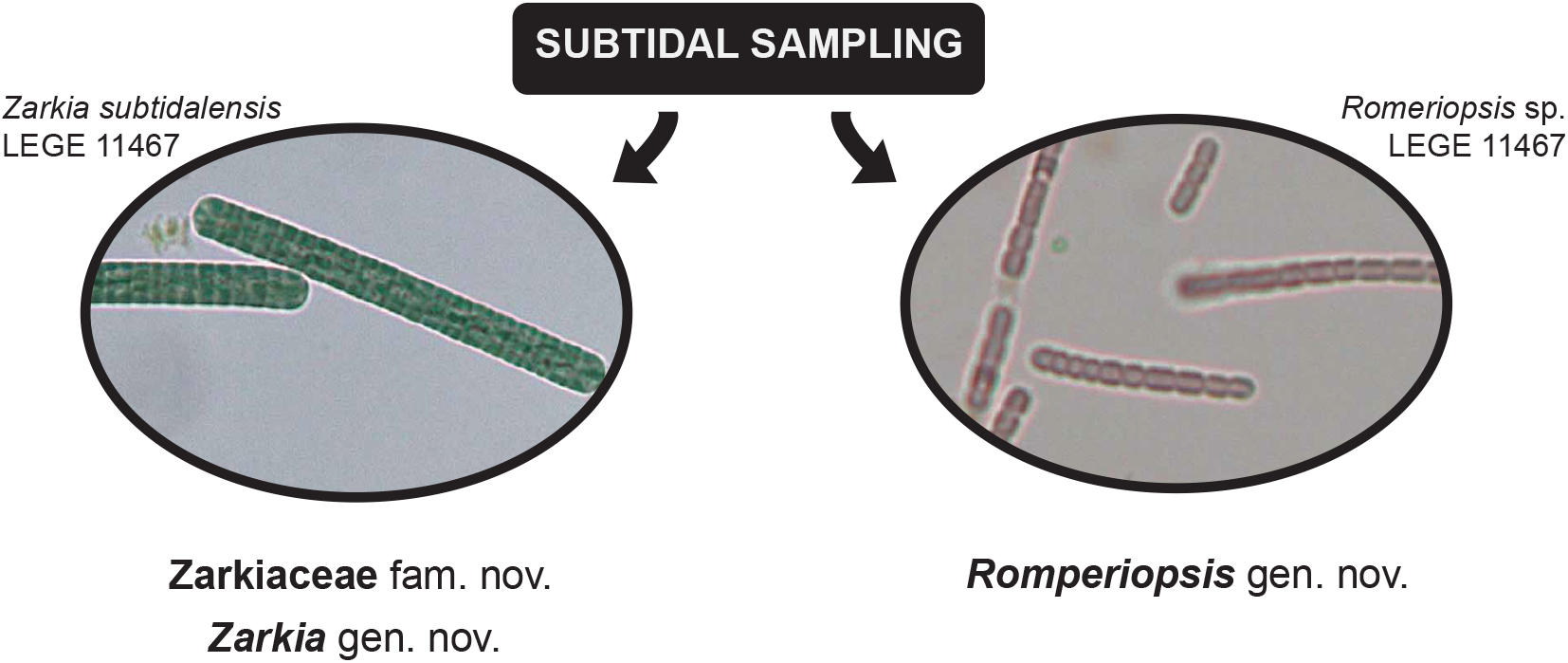

The morphology, 16S rRNA gene phylogeny and the 16S-23S rRNA gene ITS secondary structures of three strains of marine Cyanobacteria, isolated from inter- and subtidal environments from north Portugal were studied, resulting in the description of *Zarkia subtidalensis* gen. et. sp. nov. (Zarkiaceae fam. nov.) and *Romeriopsis marina* gen. et. sp. nov (Leptolyngbyaceae). No diacritical morphological characters were found either for the new family or for the new genera. The 16S rRNA gene Maximum Likelihood and Bayesian phylogenies supported that *Zarkia* and Zarkiaceae are members of the Oscillatoriales, positioned close to Microcoleaceae genera, but distant from *Microcoleus. Romeriopsis* is positioned within the Leptolyngbyaceae and is closely related to *Alkalinema*. The secondary structures of the D1-D1’, Box B, V2 and V3 helices corroborate with the phylogenetic results. Furthermore, our study supports previous observations of polyphyletic Oscillatoriales families and reinforces the need for their taxonomical revision.

## 1. Introduction

Cyanobacteria are important primary producers in the world’s oceans and shape both planktonic and benthic marine communities (Hamilton et al., 2016). In addition, they produce a plethora of secondary metabolites, such as alkaloids, polyketides, and peptides with a great variety of biological activities with some being currently used to treat cancer in the clinic (Leão et al., 2012; Calteau et al., 2014). Despite their global and biotechnological importance, the diversity of marine Cyanobacteria is still underestimated, and recently, many new genera of Cyanobacteria have been described, such as *Leptothoe* Konstantinou et Gkelis, *Marileptolyngbya* Zhou et Ling, *Salileptolyngbya* Zhou, *Lusitaniella* Ramos et al., *Dapis* Engene et al., *Capillus* and *Neolyngbya*, for example (Brito et al., 2017; Caires et al., 2018a; 2018b; Engene et al., 2018; Zhou et al., 2018; Konstantinou et al., 2019).

The growing number of new taxa descriptions, the availability of 16S rRNA gene sequences for the respective type species, and the access to computational tools that allow to expeditiously perform extensive phylogenetic reconstructions (Miller et al., 2015), has brought to light that many cyanobacterial families from the classical, morphological-based taxonomy are polyphyletic, warranting taxonomical revisions (Jahodarová et al., 2017; Nowicka-Krawczyk et al., 2018; Mai et al., 2018). Up to now, the only revision at the order level for Cyanobacteria, on the basis of robust phylogenetic analysis and morphological descriptions, concerned the Synechococcacales and was carried out by Mai et al. (2018). In that paper, the authors describe two new families and six new genera containing 14 species. The authors consider five families in the order, one of which (Trichocoleaceae) comprising a single genus. The description of families with a single genus based on a single-gene phylogenetic analysis is not a novelty (Hentschke et al., 2016) and is necessary for monophyletic clades (Johansen and Casamatta, 2005), which are well supported and not clustered in any already known family.

Against this backdrop, in this paper we describe the new family Zarkiaceae to hold the new cyanobacterial genus *Zarkia*, and also describe the new genus *Romeriopsis* (Leptolyngbyaceae), from intertidal and subtidal environments sampled in north Portugal.

## 2. Materials and Methods

### 2.1. Sampling and sites

Two samples were obtained in the subtidal zone seafloor, at 13 m depth, by collection of rocky substrate surfaces using 50 mL sterile polypropylene syringes, by SCUBA diving at ‘A Pedra’, a diving spot in front of the São Francisco Xavier fort, ∼200 m off the coast, in the city of Porto, Portugal (41.185809 N 8.719079 W). The samples were kept in 50 mL polypropylene tubes until being processed in the laboratory at CIIMAR, Porto, Portugal. These samples led to the isolation of strains *Zarkia subtidalensis* LEGE 11467 and *Romeriopsis* sp. LEGE 11480, while strain *Romeriopsis marina* LEGE 06013 had been previously isolated from a wave-exposed rock, Praia da Foz do Arelho, Caldas da Rainha (39.43327 N 9.230275 W), Portugal, as reported in Brito et al. (2012) and was obtained from the LEGE Culture Collection (LEGE-CC) at CIIMAR, Porto, Portugal (http://lege.ciimar.up.pt) (Ramos et al., 2017). The north Portuguese coastal environments, a temperate climate region, where the studied strains were collected are vulnerable to wave action, being under a strong tide and wave regime especially in the winter (Coelho et al., 2009).

### 2.2. Isolation strategy

After arrival at the laboratory, subtidal samples were observed under a light microscope. The environmental samples were not clearly dominated by cyanobacteria and therefore they were inoculated in liquid medium, supplemented with 25 g L^-1^ sea salts (Tropic Marine) and 10 µg ml^-1^ vitamin B_12_. The enrichment cultures were kept under low light conditions <10 µmol m^-2^ s^-1^ under a 14 hour light/ 10 hour dark regimen and at 19 °C. As soon as consistent growth of cyanobacteria was detected, aliquots were transferred onto solid Z8 medium plates with 1.5% agarose, supplemented with sea salts and vitamin B_12_ as described above. Liquid and solid cultures were grown at 25 °C, under a 14 hour light (approximately 30 – 40 µmol m^-2^ s^-1^)/ 10 hour dark regimen.

When single colonies or filaments were detected, these were picked with an inoculating loop and streaked onto a new medium plate. The streak plate technique was repeated until unicyanobacterial cultures were obtained following inoculation in liquid medium. The resulting unicyanobacterial strains have been deposited and since been kept in LEGE-CC under controlled temperature between 19°C and 21°C, photoperiod 14h light/10h dark, and light intensity 15–25 μmol photons m^-2^ s^-1^.

### 2.3. Morphological analysis

The morphological plasticity and cell measurements (n = 50 cells) of 50 individuals of *Zarkia* (LEGE 11467) and *Romeriopsis* strains (LEGE 06013, LEGE 11480) were examined using a Leica DMLB light microscope (Wetzlar, Germany) and micrographs were acquired with an Olympus DP73 camera and the Leica Application Suite V.4 software. The morphological characterization was made according to Komárek and Anagnostidis (2005), observing the following characters: 1) macroscopic aspect of the culture; 2) number of cells in trichomes; 3) trichome curvature and constriction; 4) shape of cells and measurements; 5) cell content; 6) presence/absence of aerotopes; 7) presence/absence of filaments sheaths or mucilage (using China Ink); 8) reproduction. The measurements were tabulated and the cells length/width were calculated for each of the new genera.

### 2.4. DNA extraction, PCR amplification and sequencing

Total genomic DNA of the three studied strains (LEGE 06013, LEGE 11480 and LEGE 11467) was isolated using MOBIO Ultraclean DNA Isolation Kit (Life Technologies). To obtain the 16S rRNA gene and the 16S-23S Internal Transcribed Spacer (ITS) of the strains LEGE 06013 and LEGE 11467, the PCR was performed using the primers 27F1 (Neilan et al., 1997) and 23SR (Taton et al., 2003) in a Biometra 2 thermal cycler (Analytik Jena). The reaction contained 13 μl H_2_O, 5 μl 5× Buffer (Promega), 2 μl MgCl_2_ (25mM), 1 μl DNTPs (10 μM), 1.25 μl of each primer (10 μM), 0.3 μl of GoTaq polymerase (Promega). The thermal cycling conditions used were: initial denaturing 94°C (5 min) followed by 10 cycles of 94°C (45 s), 57°C (45 s), 72°C (2 min), then 25 cycles of 92°C (45 s), 54°C (45 s), 72°C (2 min) before a final elongation step of 72°C (7min). The PCR products were cloned using the pGEM®–T Easy Vector System (Promega, Madison, WI, USA), transformed by heat–shock into *E. coli* cells and plated for blue–white screening (Sambrook & Russel 2001). Two colonies were selected for each strain. After growth, plasmids were extracted from white colonies using the NZYTech Miniprep Kit (NZYTech), and were prepared for sequencing using the primers 27F (Neilan et al., 1997), 359F (Nubel et al., 1997), 781R (Nubel et al., 1997), 1114F (Lane, 1991) and 23S30R (Neilan et al,. 1997). The resulting sequences were assembled using Geneious 8.1.9. software package (Biomatters) and analyzed for the presence of chimeras hidden in the rRNA sequences by DECIPHER web tool (Wright et al., 2012).

To obtain the sequence of the 16S rRNA gene of strain LEGE 11480, this gene was amplified from its gDNA using two set of primers: 27F/781R and 359F/1494R (Neilan et. al, 1997; Nubel et al., 1997) in a MyCycler (Bio-Rad laboratories Inc., Hercules, CA, USA) or T-Professional Standard (Analytik Jena) thermal cyclers, following the methodology previously described (Tamagnini et al., 1997). For this set of primers, the PCR reaction contained 7.9 μl H_2_O, 4 μl 5× Buffer (Promega), 2 μl MgCl_2_ (25mM), 1 μl DNTPs (10 μM), 2 μl of each primer (10 μM), 0.1 μl GoTaq polymerase (Promega) and 1 μl of template DNA. The thermal cycling conditions were: initial denaturation at 95 °C for 2 min, followed by 35 cycles of denaturation at 95 °C for 1 min, annealing at 55 °C for 45 s, and extension at 72 °C for 1 min, and a final extension step at 72 °C for 5 min. After PCR, to obtain the sequences, the PCR products were purified using NucleoSpin® Gel and PCR Clean-up kit (Macherey-Nagel, Düren, Germany). Purified PCR products were cloned into pGEM®-T Easy Vector (Promega, Madison, WI, USA), and transformed into OneShot TOP10 chemically competent *Escherichia coli* cells (Invitrogen, Carlsbad, CA) by heat-shock. After blue-white screening, plasmid DNA was isolated using GenElute ™ Plasmid Miniprep Kit (Sigma-Aldrich) and sequenced using the M13 primers: reverse (22mer): 5’-d

(TCACACAGGAAACAGCTATGAC)-3’ and forward (24mer): 5’-d

(CGCCAGGGTTTTCCCAGTCACGAC)-3’. The resulting sequences were assembled using Geneious 7.0. software package (Biomatters) and analyzed for the presence of chimeras hidden in the rRNA sequences by DECIPHER web tool (Wright et al., 2012).

### 2.5. 16S rRNA gene phylogenetic analysis and 16S-23S rRNA intergenic spacer (ITS) secondary structures

Phylogenies were constructed aligning *Zarkia* and *Romeriopsis* 16S rRNA gene sequences with a set of 85 sequences of homocytous cyanobacterial strains retrieved from the GenBank using BLAST. Also, we added reference strains of Synechococcales and Oscillatoriales genera and families. The total alignment length had 2384 nucleotide positions with 936 informative sites. Then, to find the phylogenetic positions of our sequences, we performed Maximum Likelihood (ML) and Bayesian Inference (BA) analysis. A similarity matrix (p-distance) was also generated to compare taxa, using MEGA version 6 (Tamura et al., 2013).

All the alignments were performed using ClustalW (Thompson et al., 1994); ML trees were performed using RAxML-HPC2 on XSEDE 8.2.10 (Stamatakis, 2014) with bootstrap = 1000; BA tree was performed using MrBayes on XSEDE 3.2.6 (Ronquist et al., 2012), with two runs of 5×10^7^ generations and NST = 6. The other parameters were left as defaults. The standard deviation for split frequencies was 0.01. All of those were ran on CIPRES Science Gateway (Miller et al., 2015).

The secondary structures of the D1-D1’, Box B, V2 and V3 helices of 16S-23S rRNA ITS were folded using Mfold (Zuker, 2003) using default parameters. The V2 helices were not found in *Pantanalinema* Vaz et al. sequences, because these ITS sequences do not present tRNAs.

## 3. RESULTS

### 3.1. Phylogenetic analysis

The BA (Fig. 1) and ML (Fig. S1) phylogenies (91 OTUs, 936 informative sites) show very identical topologies and strong statistical support in the backbones. Both trees show Oscillatoriales (BA 0.99, ML 78) and Synechococcales (BA 1, ML 37) as monophyletic orders, although Pseudanabaenaceae (traditionally Synechochoccales) is positioned at the base of both trees, outside of Oscillatoriales or Synechococcales.

**Fig. 1.**
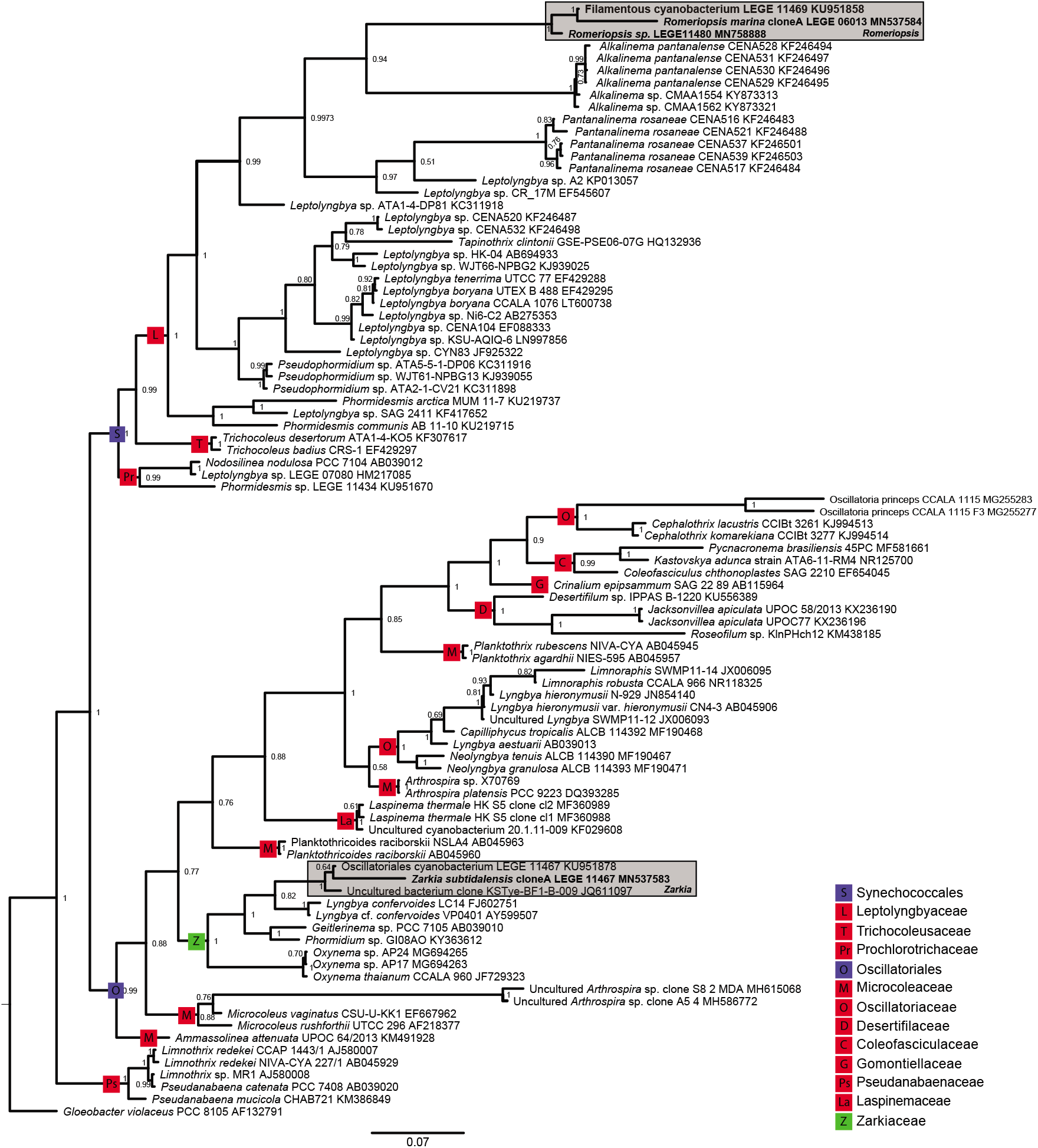
Bayesian 16S rRNA gene phylogeny constructed with trees 91 OTUs and 936 informative sites. Nodes support are presenting BA posterior probabilities. The strains sequenced for this study are in bold. Blue squares represent order level nodes. Red squares represent family level nodes.

The phylogenies confirm *Romeriopsis* as a monophyletic clade (ML=100, BA=1) in family Leptolyngbyaceae (Synechococcales), being closely related to *Alkalinema*. The new genus *Zarkia* is also monophyletic, with strong phylogenetic support (ML=100, BA=1), and is clustered with *Lyngbya cf. confervoides* (ML=79, BA=0.8), “*Phormidium*” GI08AO and *Oxynema* clades in the Oscillatoriales. In these phylogenetic trees, the order Synechococcales presents well-supported monophyletic families. The order Oscillatoriales presents polyphyletic families, such as Microcoleaceae and Oscillatoriaceae.

Although *Zarkia* is clustered with the Microcoleaceae genus *Oxynema* (ML=72, BA=1), the new genus is phylogenetically distant to *Microcoleus* and cannot be included in Microcoleaceae. Furthermore, *Zarkia* is not clustered with any other reference strain of any already known family (e. g. *Oscillatoria princeps* CCALA 115 for Oscillatoriaceae). These observations, taken together with the polyphyletic Oscillatoriales families, preclude the inclusion of the new genus *Zarkia* in any previously described family. Consequently, we describe below the new monophyletic family Zarkiaceae to encompass *Zarkia, Oxynema* Chatchawan and related clades.

The similarity matrix (p-distance) (Table S1) corroborates the phylogenetic inferences and confirms *Romeriopsis* and *Zarkia* as new genera. This analysis shows *Romeriopsis* with 98.4-99.7% of intra-clade similarity and only 89.6% of similarity to *Alkalinema*, its phylogenetically most closely-related clade. The similarity between *Zarkia* sequences is 99.1% and the similarity between this clade and *Lyngbya confervoides*, its most closely-related clade, is only 86.3%. The similarity between *Zarkia* and the clade of *Phormidium* GI08AO and *Geitlerinema* sp. PCC 7105 ranges between 93.6-94.5%.

### 3.2. 16S-23S rRNA intergenic spacer (ITS) analysis

#### 3.2.1. Zarkia

We compared the ITS secondary structure of *Zarkia* with that of its phylogenetically closest related described genus with available ITS data, *Oxynema* (Fig. 2). It was possible to compare only the D1-D1’ helix, because of the short length of the *Oxynema* CCALA 960 JF729323 sequence. Between both genera, this helix is variable in sequence, length and structure. *Zarkia* D1-D1’ helix presents a long basal stem with 10 bp plus two residues, while *Oxynema* presents the typical 4 bp (^5’^GACC^3’^ /^5’^GGTC^3’^) D1-D1’ cyanobacterial basal stem. The long basal stem of *Zarkia* is unique among Cyanobacteria. Furthermore, the 5’ side of *Zarkia*’s molecule presents no residues opposing the first lateral bulge, while *Oxynema* presents three residues in this position. These differences support the separation of both genera, which is in line with our phylogenetic analysis. It was not possible to compare *Zarkia* with *Lyngbya confervoides*, “*Phormidium*” GI08AO and “*Geitlerinema*” PCC7105, because of the lack of ITS data for these strains.

**Fig. 2.**
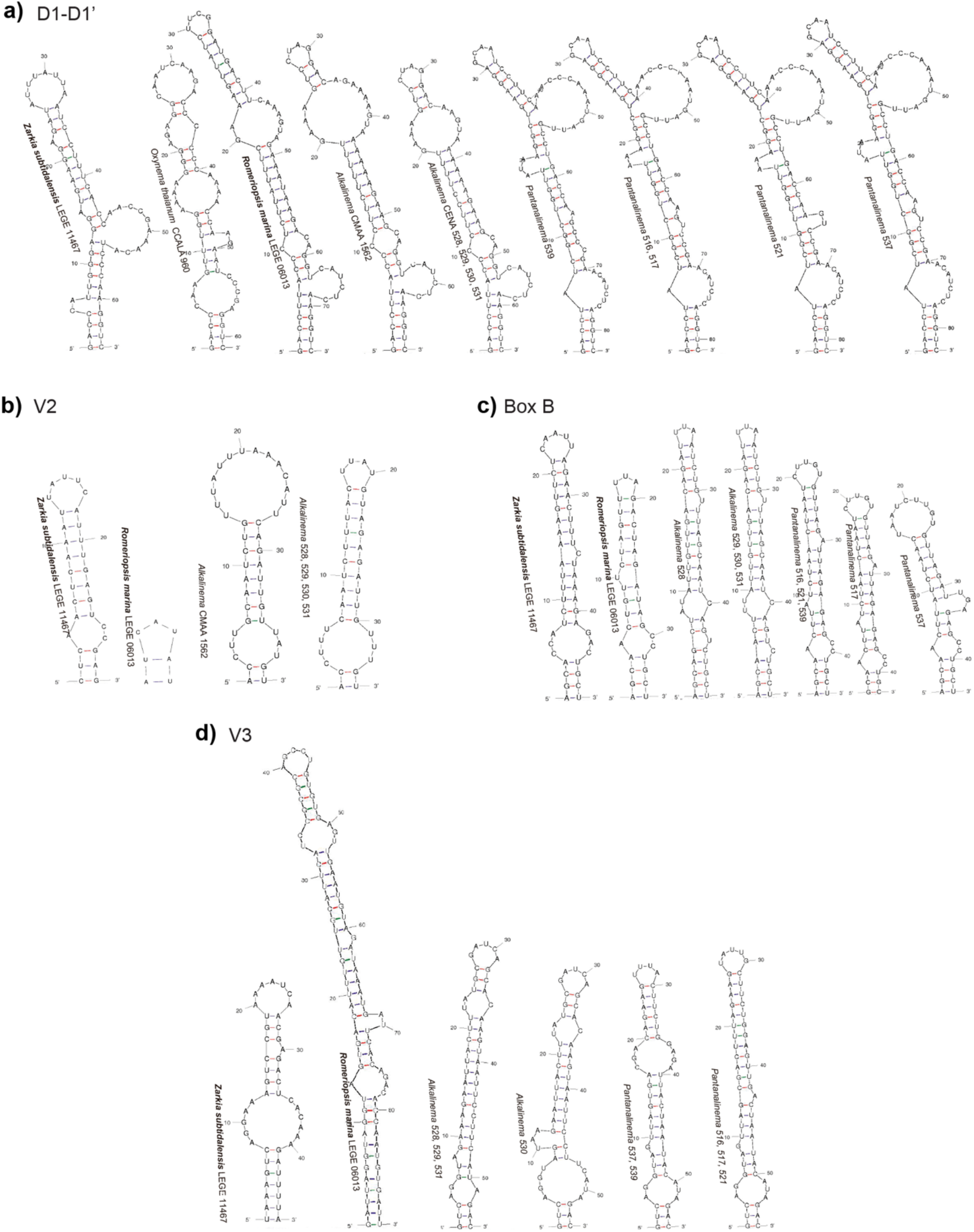
16S-23S rRNA gene ITS secondary structures of a) D1-D1’ helices, b) V2 helices, c) Box B helices and d) V3 helices of *Zarkia, Romeriopsis* and phylogenetically related genera.

#### 3.2.2. Romeriopsis

We compared the ITS secondary structures of *Romeriopsis* with those of its phylogenetically closest related genera, *Alkalinema* and *Pantanalinema* (Fig. 2). *Pantalinema*’s ITS sequences do not present the tRNAs, and because of that, the comparisons among the V2 helices were not possible with this genus. The D1-D1’ helix of *Romeriopsis* and *Alkalinema* are identical in the basal stem, lateral bulge and first loop, making the differentiation between genera impossible by this helix region. It is possible to differentiate both genera by the terminal loop. Although *Alkalinema*’s D1-D1’ helices are variable in the genus, the terminal loop is conserved (^5’^CUAG^3’^) and different from *Romeriopsis* (^5’^UUCG^3’^). The Box B, V2 and V3 helices are very different in length, sequence and structure, supporting the separation of both genera.

Comparing *Romeriopsis* with *Pantanalinema*, the D1-D1’ helix presents small variations among the strains of the latter genus. Even so, these helices are different from *Romeriopsis* helices by length, sequence and structure, remarkably by the presence of an adenine opposing the first lateral bulge in *Pantanalinema* and absence of residues in this position in *Romeriopsis*. The Box B, V2 and V3 helices are very different in length, sequence and structure, also corroborating the separation of both genera.

### 3.3. Descriptions of the new taxa

**Oscillatoriales** Schaffner

**Zarkiaceae** fam. nov. G.S. Hentschke & P. N. Leão

Filaments isopolar, solitary or entangled. Sheaths present, firm or diffluent, homogenous, opened at the ends, hyaline. Trichomes straight or wavy, cylindrical or narrowed at the ends, constricted or not constricted at cross-walls, facultatively motile. Cells isodiametric, longer than wide, shorter than wide, or discoid. Cell content sometimes granulated, with or without aerotopes. Reproduction by hormogonia.

Type: *Zarkia subtidalensis* G.S. Hentschke, A. Pinheiro, V. Ramos & P. N. Leão

***Zarkia subtidalensis*** gen. et sp. nov. G.S. Hentschke, A. Pinheiro, V. Ramos & P. N. Leão

Fig. 3

**Fig. 3.**
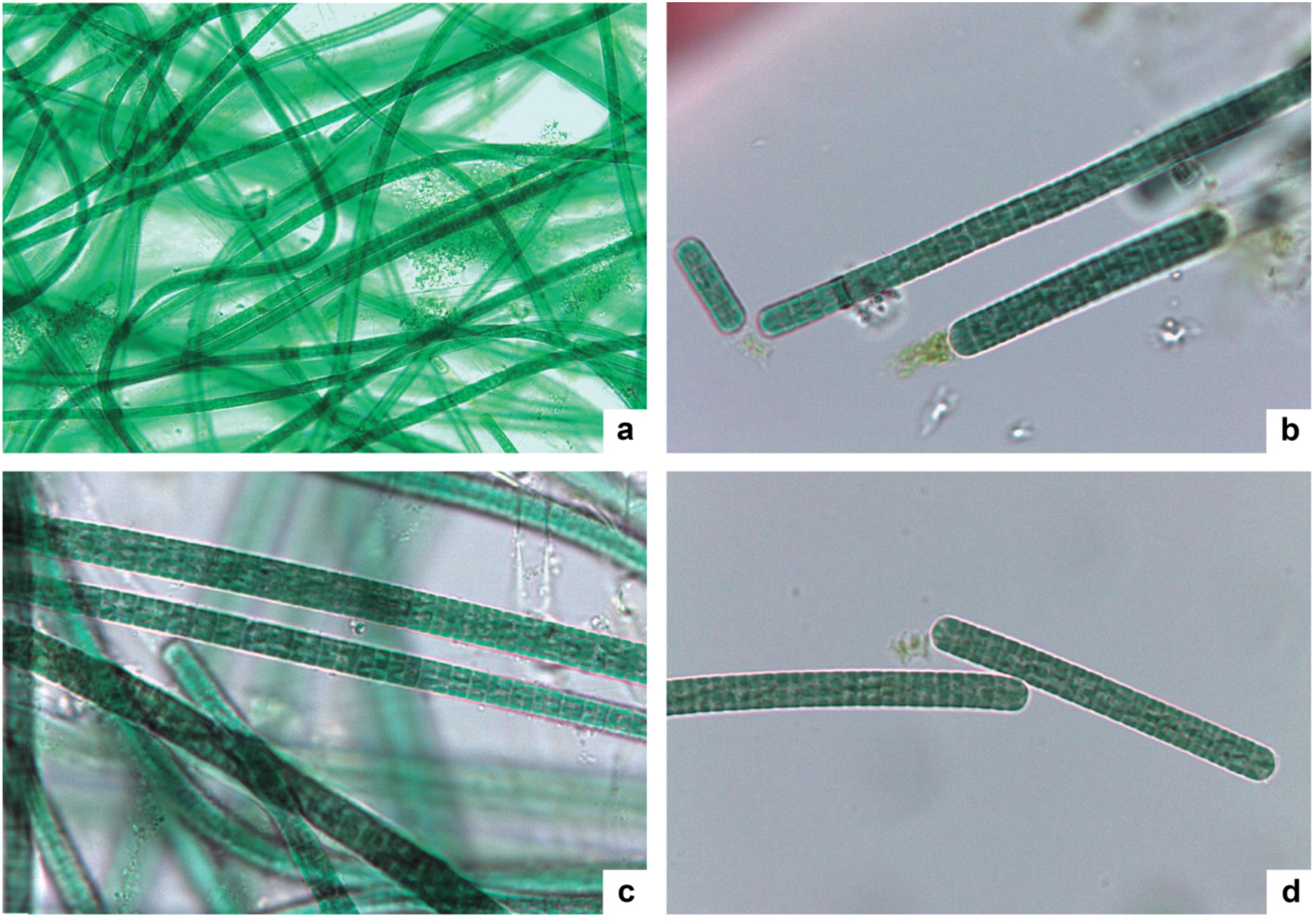
Microphotographs of *Zarkia subtidalensis*. **a**. General view of the culture; **b-d**. Details of hormogonia and trichomes showing shorter than wide cells. Magnification: **3a** = 40×, **3b-d** = 1000×.

In culture, forming mats attached to the flask walls. Filaments isopolar, solitary or entangled, straight or wavy. Sheaths firm, thin, homogenous and colourless. Trichomes cylindrical, not constricted or slightly constricted (shorter cells) at cross-walls, sometimes motile. Cells shorter than wide, rarely isodiametric or, rarely discoid (only after division), 2.7-4 μm length, 5.6-7.6 μm wide, ratio l/w 1.4-2.3 for adult cells. Apical cells rounded. Cell content homogenous, dark green, with aerotopes.

**Holotype**: PO-T4766 (unialgal population preserved lyophilized), University of Porto Herbarium.

**Type locality:** ‘A Pedra’, diving spot in front of the fort ‘Castelo do Queijo’, Portugal: (41.185809 N 8.719079 W)

**Habitat:** marine, subtidal sample, epilithic (13 m depth), about 200 m off the shore. **Etymology:** *Zarkia*, from Arabic-hispanic “zarco” means blue, the color of the ocean, and for Gonçalves Zarco Square, location of the fort near the collection site; *subtidalensis* is for subtidal.

**Reference strain:** *Zarkia subtidalensis* LEGE 11467 (MN537583) Synechococcales Hoffmann et al.

**Leptolyngbyaceae** (Komárek et Anagnostidis) Komárek et al. 2014

***Romeriopsis marina*** gen. et sp. nov. G.S. Hentschke, A. Pinheiro, V. Ramos et P. N. Leão

Fig. 4a-d

**Fig. 4.**
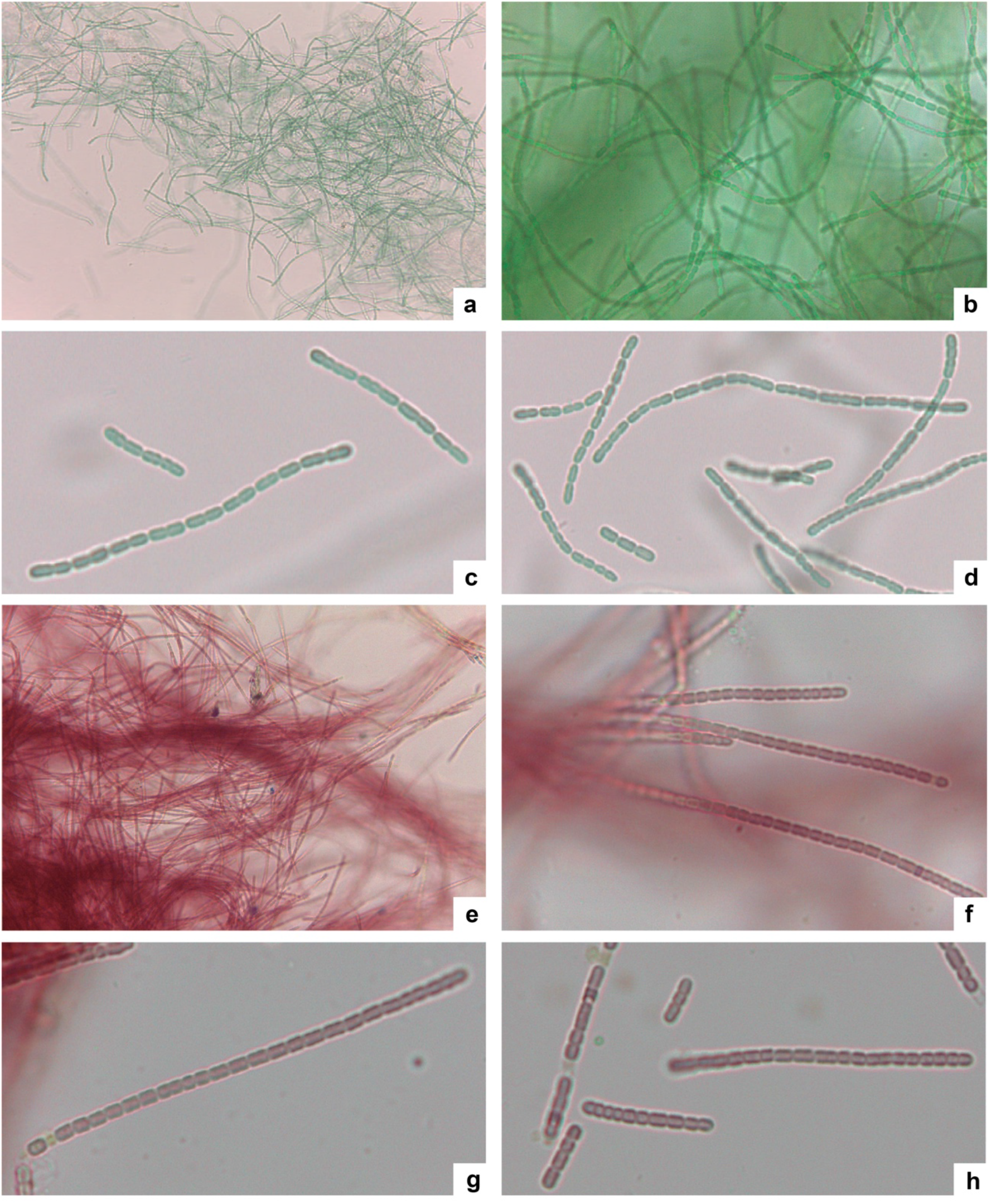
Microphotographs of *Romeriopsis marina* (**a-d**) and *Romeriopsis* sp. (**e-h**). **a, b** – General view of the culture showing long and short trichomes; **c, d** – Details of short trichomes. **e** – General view of the culture showing long and short trichomes; **f-h** – Details of short trichomes. Magnification: **4a, 4e** = 40×; **4b-d** and **4f-h** = 1000×.

In culture, trichomes solitary or forming fluffy clusters not attached to the flask walls. Trichomes cylindrical, constricted, slightly curved or wavy, few celled (more common) or long (>50 cells check other species). With very thin sheaths (visible only in broken filaments) or without. Without mucilaginous envelope. Adult cells longer than wide, cylindrical or rarely barrel shaped, 1.6-3 μm length, 1.5-2 μm wide, ratio length/width 1.2-1.9. Terminal cells rounded. Cell content olive-green, not granulated, without aerotopes. Reproduction by fragmentation of trichomes.

**Holotype:** PO-T4767 (unialgal population preserved lyophilized), University of Porto Herbarium.

**Type locality:** Praia da Foz do Arelho, Caldas da Rainha, Portugal, (39.43327 N 9.230275 W)

**Habitat:** marine, intertidal zone, wave-exposed rock

**Etymology:** *Romeriospsis*, similar to *Romeria*, due to the short trichomes; *marina* is for marine.

**Reference strain:** *Romeriopsis marina* LEGE 06013 MN537584

Notes

Differs from *Romeria* by the common presence of long trichomes (>50 cells), which do not disintegrate easily, by the presence of sheaths and by the absence of diffluent mucilaginous envelopes. Indistinguishable in morphology from *Pantanalinema* and *Alkalinema*.

*Romeriopsis* sp. LEGE 11480 (Fig. 4, e-h) was identified only in generic level. This strain is morphologically identical to *Romeriospsis marina* LEGE 06013 but presents reddish cell content and was collected in a different habitat. *Romeriopsis* sp. LEGE 11480 was collected at the same location and habitat as *Zarkia subtidalensis* LEGE 11467, on a rocky substrate at 13m depth and 200 m off the shore, being subtidal, while *R. marina* LEGE 06013 was collected on a wave-exposed rock in the intertidal zone. Although the 16S rRNA gene tree indicates that *Romeriopsis* sp. LEGE 11480 could be another species in *Romeriopsis* clade, we could not describe it, because of the lack of ITS data and any morphological diagnostic character.

## 4. Discussion

The wave energy action and related environmental conditions from the north continental Portuguese coast (Coelho et al., 2009), where the studied marine strains were collected, are known drivers shaping the coastal biodiversity, including that of microbial communities (Witt et al., 2012). Along with the sampling effort, this may partially explain why this marine temperate region is being a rich source of novel cyanobacteria taxa, already described (Brito et al., 2017) or at least phylogenetically highlighted (Ramos et al., 2018).

In this paper we described the new genera *Romeriopsis* and *Zarkia*, and the new family Zarkiaceae supported by 16S rRNA gene phylogeny and 16S-23S rRNA ITS secondary structures. The family Zarkiaceae was named after *Zarkia* and encompasses a monophyletic clade containing *Zarkia*, as well as the strains *Lyngbya confervoides* LC14 FJ602751, *Lyngbya* cf. *confervoides* VP0401 AY599507, *Phormidim* sp. GI08AO KY363612, *Geitlerinema* sp. PCC 7105 AB039010 and the genus *Oxynema*. The strains identified as *Lyngbya confervoides*, were previously used as reference for *Lyngbya* by Komárek et al. (2013) and Caires et al. (2018a), but although they are morphologically and ecologically similar to the type species description, they are not well characterized (Komárek et al., 2013) and cannot be surely assigned to this genus, because of limited available data (only one photo and not conclusive phylogeny in Sharp et al., 2009). Many other papers show *Lyngbya* as a polyphyletic genus (Engene et al., 2018; Caires et al., 2018b; Mühlsteinová et al., 2018) and to identify the true *Lyngbya* clade, a robust revision of the genus is needed. This has been done, for example, by Mühlsteinová et al. (2018) to establish the true *Oscillatoria* clade. According to this, we didn’
st include the genus *Lyngbya* in Zarkiaceae and chose to consider the *Lyngbya* cf. *confervoides* clade as still undefined, worthy to be revised., The monophyletic clade containing “*Phormidium*” GI08AO and “*Geitlerinema*” PCC 7105 strains is another genus that has to be described in the future.

Morphologically, Zarkiaceae is similar to Microcoleaeceae, presenting, in the case of *Zarkia*, isodiametric or slightly shorter than wide adult cells, and is it also similar to Oscilatoriaceae presenting discoid cells, in the case of *Lyngbya* cf. *confervoides* strains. As for the descriptions of Ocullatelaceae and Trichocoleaceae (Synechococcales) (Mai et al., 2018), no diacritical morphological markers were found for Zarkiaceae. Although this lack of morphological diacritic characters, according to our trees, we could not assign *Zarkia* to any already described family, considering that this genus is phylogenetically unrelated to any reference strains for previously-described families represented here by *Desertifilum* sp. IPPAS B-1220 KU556389 (Desertifilaceae), *Coleofasciculus chthonoplastes* SAG 2210 EF654045 (Coleofasciculaceae), *Crinalium epipsammum* SAG 22.89 NR_112218 (Gomontiellaceae), *Microcoleus vaginatus* CSU-U-KK1 EF66796 (Microcoleaceae) and *Oscillatoria princeps* CCALA 1115 F3 MG255277 (Oscillatoriaceae). Furthermore, it is evident that Oscillatoriales families are polyphyletic. Our trees show six clades containing Microcoleaceae genera, five of which not related to *Microcoleus*. Likewise, there is one additional Oscillatoriaceae clade (*Capilliphycus* and *Neolyngbya* clade) not related to *Oscillatoria*. Although we find Oscillatoriales as a monophyletic order, our phylogenies are in agreement with Ishida et al. (2001), Jahodarová et al. (2017) and Nowicka-Krawczyk et al. (2018), which also show many polyphyletic Oscillatoriales families, including Microcoleaceae and Oscillatoriaceae. We highlight also that the family Pseudanabaenaceae (traditionally Synechococcales) is not included in Oscillatoriales or Synechococcales clades, indicating that the phylogenetic position and taxonomy of this family must be revised. The same result was found by Mai et al (2018).

In this study we use the monophyletic species concept (Johansen and Casamatta, 2005) for delimitation of genera and families (Komárek et al., 2014; Mai et al., 2018). As stated by Komárek et al. (2014), “morphological characters used to define higher taxa…have apparently arisen and/or been lost several times during the evolution of modern species and genera”. Considering this, we believe that in the near future, after taxonomical revisions based on 16S rRNA gene, cryptic cyanobacterial families will be more common. Mai et al. (2018) highlight the importance of finding new markers for higher taxonomical levels; in order to describe Ocullatelaceae and Trichocoleaceae, they have used molecular markers in the 16S rRNA gene secondary structures and *rpco1* gene phylogenies, which, however were not always in agreement with the 16S rRNA gene phylogenies. Due to this, we considered the 16S rRNA gene phylogenies as the current instrument to separate families, which, combined with the monophyletic nature of Zarkiaceae, led to the description of a new family. The 16S-23S rRNA gene ITS secondary structures support our proposal, since *Zarkia* presents a unique D1-D1’ helix among Cyanobacteria.

In the Synechococcales clade, our phylogenetic analysis (Fig. 1) is in agreement with the revision of Mai et al. (2018) for the order, showing monophyletic families. *Romeriopsis* is clearly monophyletic (ML=100, BA=1) and positioned in Leptolyngbyaceae, clustered with *Alkalinema* and *Pantanalinema*. A smaller related clade contains strains assigned to *Leptolyngbya* (*Leptolyngbya* sp. A2 KP01305 and *Leptolyngbya* sp. CR_17M EF545607), which must be described as a new genus in the future. The 16S-23S rRNA gene ITS secondary structures are also in agreement with these findings, as explained in the results section.

The herein described new genera, *Zarkia* and *Romeriopsis*, are supported by 16S rRNA phylogenies, 16S-23S rRNA gene ITS secondary structures and morphological analysis. *Zarkia* is grouped with high phylogenetic support with *Lyngbya* cf. *confervoides*, but these taxa are morphologically very different. *Zarkia* presents adult cells mainly smaller than wide (ratio l/w 1.4-2.3), discoid only after division, while *Lyngbya* cf. confervoides presents the typical *Lyngbya* discoid adult cells. The 16S-23S rRNA gene ITS secondary structures of the D1-D1’ helix of *Zarkia* are unique among Cyanobacteria, as commented above. *Romeriopsis* is morphologically similar to *Romeria* regarding a generally short trichome length but differs by the common presence of long trichomes (>50 cells), which do not disintegrate easily as in *Romeria*, by the presence of sheaths and the absence of diffluent mucilaginous envelopes. Up to now, there is no available 16S rRNA gene sequence clearly assigned to *Romeria* in public databases. For this reason, we have not included this genus in our phylogeny. *Romeriospsis* is morphologically indistinguishable from *Pantanalinema* and *Alkalinema*, but the phylogenies and the 16S-23S rRNA gene ITS secondary structures in the current study confirm that these are three separate genera.

## 5. Conclusion

The 16S rRNA gene is still the most reliable molecular marker for taxonomical studies at the genus level and is proving to be useful also for family delimitation of Cyanobacteria, although additional markers must be tested. For that reason, a global effort from the research community is required to allow the reconstruction of the evolutionary history of these photosynthetic microorganisms. Traditional, morphological-based family taxonomy is in conflict with present molecular and phylogenetic data. By proposing the erection of a new family, this work is an additional contribution towards the long journey of resolving Cyanobacteria systematics. Based on the phylogenetic data presented herein, we also emphasize the need for a taxonomic revision of several Oscillatoriales families and genera.

## Statement of informed consent

No conflicts, informed consent, human or animal rights applicable.

## Declaration of the contributions of the authors

GSH worked in collection and assembly of data, analysis and interpretation of the data and drafting the article. AP, VR, AB, MSC, SB, VV, PNL worked in conception and design, collection and assembly of data, obtaining of funding, analysis and interpretation of the data and critical revision of the article for important intellectual content. All authors read and approved the final version of the manuscript.

## Supporting Information

**Fig. S1**. Maximum Likelihood 16S rRNA gene phylogeny performed with 91 OTUs and 936 informative sites.

**Table S1**. Similarity matrix (p-distance) comparing the 16S rRNA gene of *Zarkia, Romeriopsis* and related strains.

## References

Brito, Â., Ramos, V., Mota R., Lima S., Santos, A., Vieira, J., Vieira, C. P., Kaštovsky, J., Vasconcelos, V. M., Tamagnini, P., 2017. Description of new genera and species of marine cyanobacteria from the Portuguese Atlantic coast. Mol. Phyl. Evol. 111, 18–34. https://doi.org/10.1016/j.ympev.2017.03.006.

Caires, T. A., Lyra, G. M., Hentschke, G. S., da Silva, A. M. S., de Araújo, V. L., Sant’Anna, C. L., Nunes J. M. C., 2018a. Polyphasic delimitation of a filamentous marine genus, Capillus gen. nov. (Cyanobacteria, Oscillatoriaceae) with the description of two Brazilian species. Algae 33(4), 291–304. https://doi.org/10.4490/algae.2018.33.11.25

Caires, T. A., Lyra, G. M., Hentschke, G. S., Pedrini, A. G., Sant’Anna, C. L., Nunes, J. M. C., 2018b. Neolyngbya gen. nov. (Cyanobacteria, Oscillatoriaceae): A new filamentous benthic marine taxon widely distributed along the Brazilian coast. Mol. Phyl. Evol. 120, 196–211. https://doi.org/10.1016/j.ympev.2017.12.009

Calteau, A., Fewer, D. P., Latifi, A., Coursin, T., Laurent, T., Jokela, J., Kerfeld, C. A., Sivonen, K., Piel, J., Gugger, M., 2014. Phylum-wide comparative genomics unravel the diversity of secondary metabolism in Cyanobacteria. BMC Genomics, 15(1), 977. https://doi.org/10.1186/1471-2164-15-977

Coelho, C., Silva, R., Veloso-Gomes, F., Taveira-Pinto, F., 2009. Potential effects of climate change on northwest Portuguese coastal zones. ICES Journal of Marine Science 667, 1497–1507.

Engene, N., Tronholm, A., Paul, V. J., 2018. Uncovering cryptic diversity of Lyngbya: the new tropical marine cyanobacterial genus Dapis (Oscillatoriales). J. Phycol. 54(4), 435–446. https://doi.org/10.1111/jpy.12752.

Hamilton, T. L., Bryant, D. A., Macalady, J. L., 2016. The role of biology in planetary evolution: cyanobacterial primary production in low-oxygen Proterozoic oceans. Environ. Microbiol., 18(2), 325–340. https://doi.org/10.1111/1462-2920.13118.

Hentschke, G. S., Johansen, J. R., Pietrasiak, N., Fiore, M. F., Rigonato, J., Sant’Anna, C. L., Komárek, J., 2016. Phylogenetic placement of Dapisostemon gen. nov. and Streptostemon, two tropical heterocytous genera (Cyanobacteria). Phytotaxa 245 (2), 129–143. https://doi.org/10.11646/phytotaxa.245.2.4.

Ishida, T., Watanabe, M.M., Sugiyama, J. & Yokota A., 2001. Evidence for polyphyletic origin of the members of the orders of Oscillatoriales and Pleurocapsales as determined by 16S rDNA analysis. FEMS Microbiol. Lett. 201(1), 79–82. https://doi.org/10.1111/j.1574-6968.2001.tb10736.x.

Jahodářová, E., Dvořák, P., Hašler, P., Holušová, K., Poulíčková, A., 2017. Elainella gen. nov.: a new tropical cyanobacterium characterized using a complex genomic approach. Eur. J. Phycol. 53(1), 39–51. https://doi.org/10.1080/09670262.2017.1362591

Johansen, J. R., Casamatta, D. A., 2005. Recognizing cyanobacterial diversity through adoption of a new species paradigm. Algol. Stud. 117, 71–93. https://doi.org/10.1127/1864-1318/2005/0117-0071

Komárek, J., Anagnostidis, K., 2005. Cyanoprokaryota 2. Teil Oscillatoriales, in: Büdel, B., Krienitz, L., Gärtner, G., Schagerl, M. (Eds.), Süsswasserflora von Mitteleuropa, vol. 19/2. Elsevier Spektrum Akademische, München, pp 1–759.

Komárek, J., Zapomělová, E., Šmarda, J., Kopecký, J., Rejmánková, E., Woodhouse, J., Neilan, B. N., Komárková, J., 2013. Polyphasic evaluation of Limnoraphis robusta, a water–bloom forming cyanobacterium from Lake Atitlán, Guatemala, with a description of Limnoraphis gen. nov.. Fottea 13(1), 39–52. https://doi.org/10.5507/fot.2013.004

Komárek, J., Kastovsky, J., Mares, J., Johansen, J. R., 2014. Taxonomic classification of cyanoprokaryotes (cyanobacterial genera), using a polyphasic approach, Preslia 86, 295–335. https://doi.org/10.1080/09670262.2016.1163738

Konstantinou, D., Voultsiadou, E., Panteris, E., Zervou, S., Gkelis A. H. S., 2019. Leptothoe, a new genus of marine cyanobacteria (Synechococcales) and three new species associated with sponges from the Aegean Sea. J. Phycol. 55(4), 882–897. https://doi.org/10.1111/jpy.12866.

Kotai, J., 1972. Instructions for Preparation of Modified Nutrient Solution Z8 for Algae Publication B-11/69. Norwegian Institute for Water Research, Oslo.

Lane, D. J., 1999. 16S/23S rRNA sequencing, in: Stackebrandt, E., Goodfellow M. (Eds.), Nucleic Acid Techniques in Bacterial Systematics. Wiley, Chichester, pp. 115– 175.

Mai, T., Johansen, J. R., Pietrasiak, N., Bohunická, M., Martin, M. P., 2018. Revision of the Synechococcales (Cyanobacteria) through recognition of four families including Oculatellaceae fam. nov. and Trichocoleaceae fam. nov. and six new genera containing 14 species. Phytotaxa 365(1), 1–59. https://doi.org/10.11646/phytotaxa.365.1.1.

Miller, M., Schwartz, T., Pickett, B., He, S., Klem, E., Passarotti, R. H.M., Kaufman, S., O’Leary, M. A., 2015. A RESTful API for Access to Phylogenetic Tools via the CIPRES Science Gateway. Evol. Bioinform. 11, 43–8. https://doi.org/10.4137/EBO.S21501

Mühlsteinová, R., Hauer, T., De Ley, P., Pietrasiak, N., 2018. Seeking the true Oscillatoria: A quest for a reliable phylogenetic and taxonomic reference point. Preslia 90, 151–169. https://doi.org/10.23855/preslia.2018.151.

Neilan, B. A., Jacobs, D., Dot, T. D., Blackall, L. L., Hawkins, P. R., Cox P. T., Goodman, A. E., 1997. rRNA sequences and evolutionary relationships among toxic and nontoxic cyanobacteria of the genus Microcystis. Int. Jour. Syst. Bact. 47, 693–697. https://doi.org/10.1099/00207713-47-3-693

Nowicka-Krawczyk, P., Mühlsteinová, R., Hauer, T., 2019. Detailed characterization of the Arthrospira type species separating commercially grown taxa into the new genus Limnospira (Cyanobacteria). Nat. Scient. Rep. 9, 694. https://doi.org/10.1038/s41598-018-36831-0.

Nubel, U., Garcia-Pichel, F., Muyzer, G., 1997. PCR primers to amplify 16S rRNA genes from cyanobacteria. Appl. Environ. Microbiol. 63, 3327–3332.

Ramos et al., 2017. Cyanobacterial diversity held in microbial biological resource centers as a biotechnological asset: the case study of the newly established LEGE culture collection. Journal of Applied Phycology 30, 1437–1451.

Ronquist, F., Teslenko, M., Van der Mark, P., Ayres, D. L., Darling, A., Höhna, S., Larget, B., Liu, L., Suchard, M. A., Huelsenbeck, J. P., 2012. MrBayes 3.2: Efficient bayesian phylogenetic inference and model choice across a large model space. Syst. Biol. 61, 539–542. https://doi.org/10.1093/sysbio/sys029.

Sambrook, J., Russell, D.W., 2001. Molecular Cloning, volume 1: a laboratory manual. CSHL Press, New York.

Sharp, K., Arthur, K. E., Gu, L., Ross, C., Harrison, G., Gunasekera, S. P., Meickle, T., Matthew, S., Luesch, H., Thacker, R. W., Sherman, D. H., Paul, V. J., 2009. Phylogenetic and chemical diversity of three chemotypes of bloom-forming Lyngbya species (Cyanobacteria: Oscillatoriales) from reefs of southeastern Florida. Appl. Environ. Microbiol. 75(9), 2879–2888. https://doi.org/10.1128/AEM.02656-08

Stamatakis, A., 2014. RAxML version 8: a tool for phylogenetic analysis and post-analysis of large phylogenies. Bioinf. 30, 1312–1313. https://doi.org/10.1093/bioinformatics/btu033.

Tamura, K., Stecher, G., Peterson, D., Filipski, A., Kumar, S., 2013. MEGA6: molecular evolutionary genetics analysis version 6.0. Mol. Biol. Evol. 30, 2725–2729. https://doi.org/10.1093/molbev/mst197.

Taton, A., Grubisic, S., Brambilla, E., De Wit, R., Wilmotte, A., 2003. Cyanobacterial diversity in natural and artificial microbial mats of Lake Fryxell (McMurdo Dry Valleys, Antarctica): a morphological and molecular approach. Appl. Environ. Microbiol. 69(9), 5157–5169. https://doi.org/10.1128/aem.69.9.5157-5169.

Thompson, J. D., Higgins, D. G., Gibson, T. J., 1994. CLUSTAL W: improving the sensitivity of progressive multiple sequence alignment through sequence weighting, position-specific gap penalties and weight matrix choice. Nucl. Acids Res. 22(22), 4673–4680. https://doi.org/10.1093/nar/22.22.4673.

Witt, M. J., Sheehan, E. V., Bearhop, S., Broderick, A. C., Conley, D. C., Cotterell, S. P., Godley, B. J., 2012. Assessing wave energy effects on biodiversity: the Wave Hub experience. Philosophical Transactions of the Royal Society A: Mathematical, Physical and Engineering Sciences 370, 502–529.

Wright, E. S., Yilmaz, L. S., Noguera, D. R., 2012. DECIPHER, A Search-Based Approach to Chimera Identification for 16S rRNA Sequences. Appl. Environ. Microbiol. 78(3), 717–725. https://doi.org/10.1128/AEM.06516-11.

Zhou, W. G., Ding, D. W., Yang, Q. S., Ahmad, M., Zhang, Y. Z., Lin, X. C., Zhang, Y. Y., Ling, J., Dong, J. D., 2018. Marileptolyngbya sina gen. nov., sp. nov. and Salileptolyngbya diazotrophicum gen. nov., sp. nov. (Synechococcales, Cyanobacteria), species of cyanobacteria isolated from a marine ecosystem. Phytotaxa 383, 075–092. https://doi.org/10.11646/phytotaxa.383.1.4.

Zuker, M., 2003. Mfold web server for nucleic acid folding and hybridization prediction. Nucl. Acids Res. 31, 3406–3415. https://doi.org/10.1093/nar/gkg595.

