## Supporting Information for "Establishment of the family Zarkiaceae (Oscillatoriales, Cyanobacteria) and description of the new marine genera *Zarkia* (Zarkiaceae, Oscillatoriales) and *Romeriopsis* (Leptolyngbyaceae, Synechococcales), from northern Portugal"

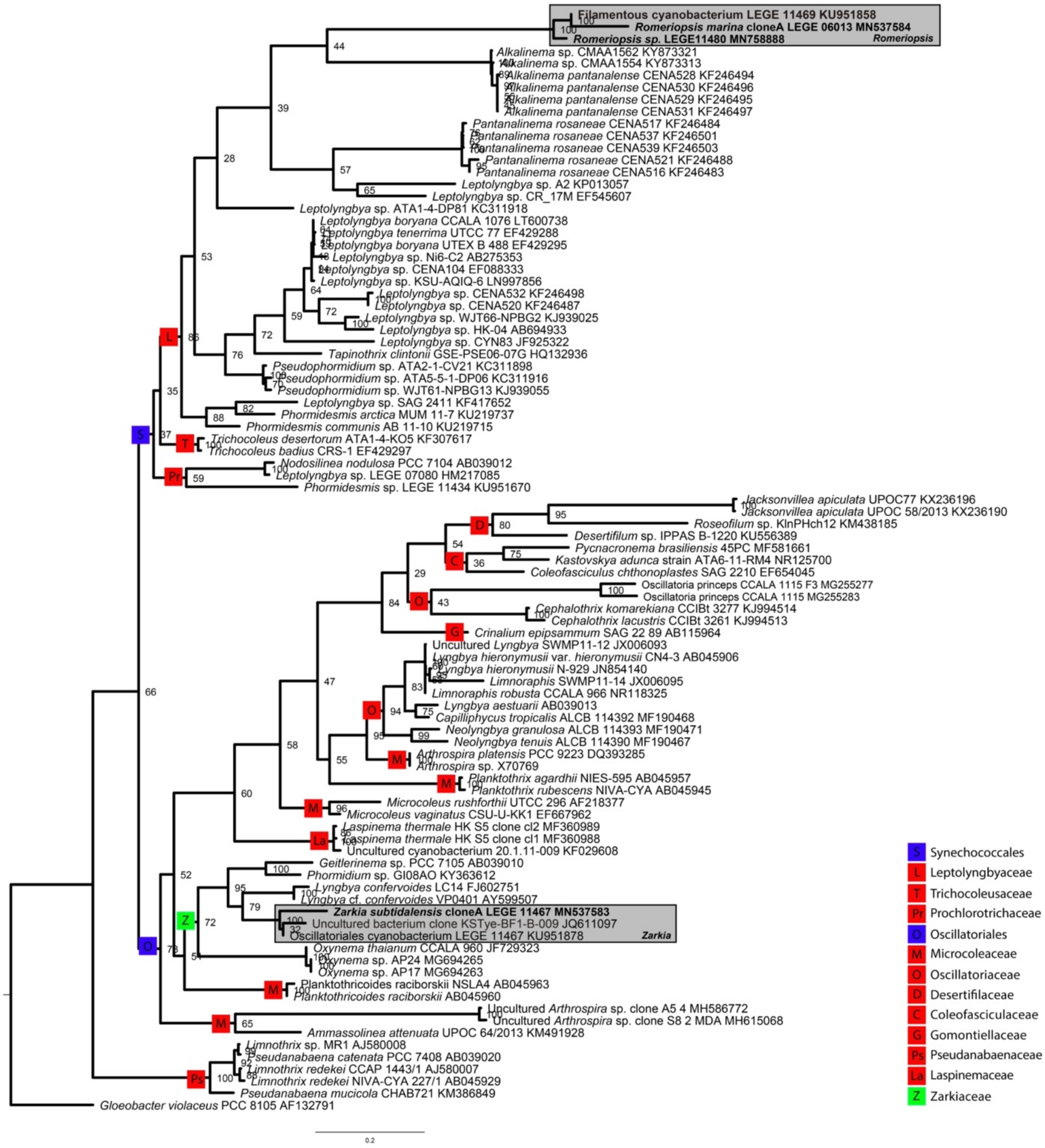


**Fig. S1.** Maximum Likelihood 16S rRNA gene phylogeny performed with 91 OTUs and 936 informative sites.

**Table S1.** Similarity matrix (p-distance) comparing the 16S rRNA gene of *Zarkia, Romeriopsis* and related strains.

| | | 1 | 2 | 3 | 4 | 5 | 6 | 7 | 8 | 9 | 10 | 11 | 12 | 13 | 14 | 15 | 16 | 17 | 18 | 19 | 20 | 21 | 22 | 23 |
|---|---|---|---|---|---|---|---|---|---|---|---|---|---|---|---|---|---|---|---|---|---|---|---|---|
| 1 | MN537584 *Romeriopsis marina* LEGE 06013 | | | | | | | | | | | | | | | | | | | | | | | |
| 2 | KU951858 Filamentous cyanobacterium LEGE11469 | 99.7 | | | | | | | | | | | | | | | | | | | | | | |
| 3 | MN758888 *Romeriopsis marina* LEGE 11480 | 98.7 | 98.6 | | | | | | | | | | | | | | | | | | | | | |
| 4 | KF246494 *Alkalinema pantanalense* CENA528 | 89.4 | 89.5 | 89.2 | | | | | | | | | | | | | | | | | | | | |
| 5 | KY873321 *Alkalinema* sp. CMAA1562 | 89.6 | 89.7 | 89.6 | 99.5 | | | | | | | | | | | | | | | | | | | |
| 6 | KY873313 *Alkalinema* sp. CMAA1554 | 89.6 | 89.7 | 89.4 | 99.5 | 99.8 | | | | | | | | | | | | | | | | | | |
| 7 | KF246497 *Alkalinema pantanalense* CENA531 | 89.6 | 89.7 | 89.5 | 99.8 | 99.7 | 99.7 | | | | | | | | | | | | | | | | | |
| 8 | KF246496 *Alkalinema pantanalense* CENA530 | 89.5 | 89.6 | 89.4 | 99.7 | 99.6 | 99.6 | 99.9 | | | | | | | | | | | | | | | | |
| 9 | KF246495 *Alkalinema pantanalense* CENA529 | 89.6 | 89.7 | 89.5 | 99.8 | 99.7 | 99.7 | 100 | 99.9 | | | | | | | | | | | | | | | |
| 10 | KP013057 *Leptolyngbya* sp. A2 | 88.8 | 88.9 | 88.4 | 88.8 | 89.3 | 89.1 | 89.1 | 89.0 | 89.1 | | | | | | | | | | | | | | |
| 11 | EF545607 *Leptolyngbya* sp. CR_17M | 88.4 | 88.8 | 88.3 | 92.4 | 92.9 | 92.7 | 92.7 | 92.6 | 92.7 | 92.0 | | | | | | | | | | | | | |
| 12 | KF246483 *Pantanalinema rosaneae* CENA516 | 88.4 | 88.5 | 87.8 | 90.9 | 91.3 | 91.3 | 91.2 | 91.1 | 91.2 | 91.1 | 92.6 | | | | | | | | | | | | |
| 13 | KF246488 *Pantanalinema rosaneae* CENA521 | 88.0 | 88.1 | 87.5 | 90.6 | 91.0 | 90.9 | 90.8 | 90.7 | 90.8 | 90.7 | 92.3 | 99.7 | | | | | | | | | | | |
| 14 | KF246501 *Pantanalinema rosaneae* CENA537 | 88.4 | 88.5 | 87.9 | 90.8 | 91.4 | 91.4 | 91.1 | 90.9 | 91.1 | 91.1 | 92.6 | 99.9 | 99.6 | | | | | | | | | | |
| 15 | KF246503 *Pantanalinema rosaneae* CENA539 | 88.3 | 88.4 | 87.8 | 90.7 | 91.3 | 91.3 | 90.9 | 90.8 | 90.9 | 91.0 | 92.5 | 99.8 | 99.5 | 99.9 | | | | | | | | | |
| 16 | KF246484 *Pantanalinema rosaneae* CENA517 | 88.3 | 88.4 | 87.8 | 90.7 | 91.3 | 91.3 | 90.9 | 90.8 | 90.9 | 91.0 | 92.5 | 99.8 | 99.5 | 99.9 | 99.8 | | | | | | | | |
| 17 | JQ611097 Uncultured bacterium KSTyeBF1B009 | 86.3 | 86.1 | 85.6 | 86.5 | 87.0 | 87.0 | 86.7 | 86.6 | 86.7 | 86.5 | 86.6 | 89.5 | 89.2 | 89.5 | 89.4 | 89.4 | | | | | | | |
| 18 | KU951878 *Zarkia subtidalensis* LEGE11467 | 85.6 | 85.5 | 85.0 | 87.4 | 87.9 | 87.9 | 87.6 | 87.5 | 87.6 | 86.7 | 87.0 | 89.1 | 88.8 | 89.1 | 89.0 | 89.0 | 99.1 | | | | | | |
| 19 | FJ602751 *Lyngbya confervoides* LC14CAIRES | 86.3 | 86.1 | 85.6 | 86.8 | 87.3 | 87.3 | 87.0 | 86.9 | 87.0 | 85.5 | 86.7 | 87.8 | 87.4 | 87.8 | 87.7 | 87.7 | 95.8 | 95.6 | | | | | |
| 20 | AY599507 *Lyngbya* cf. *confervoides* VP0401CAIRES | 86.3 | 86.1 | 85.6 | 86.8 | 87.3 | 87.3 | 87.0 | 86.9 | 87.0 | 85.5 | 86.7 | 87.8 | 87.4 | 87.8 | 87.7 | 87.7 | 95.8 | 95.6 | 100 | | | | |
| 21 | MG694265 *Oxynema* sp. AP24 | 83.5 | 83.3 | 82.6 | 86.2 | 86.4 | 86.4 | 86.4 | 86.3 | 86.4 | 85.7 | 86.1 | 87.7 | 87.4 | 87.7 | 87.6 | 87.6 | 90.3 | 90.0 | 91.0 | 91.0 | | | |
| 22 | MG694263 *Oxynema* sp. AP17 | 83.5 | 83.3 | 82.6 | 86.2 | 86.4 | 86.4 | 86.4 | 86.3 | 86.4 | 85.7 | 86.1 | 87.7 | 87.4 | 87.7 | 87.6 | 87.6 | 90.3 | 90.0 | 91.0 | 91.0 | 100 | | |
| 23 | KY363612 *Phormidium* sp. GI08AO | 86.2 | 85.8 | 85.6 | 87.1 | 87.7 | 87.6 | 87.4 | 87.3 | 87.4 | 84.9 | 86.0 | 87.7 | 87.3 | 87.7 | 87.5 | 87.5 | 94.0 | 94.5 | 93.3 | 93.3 | 89.8 | 89.8 | |
| 24 | AB039010 *Geitlerinema* sp. PCC7105 | 83.5 | 83.4 | 83.1 | 86.5 | 86.7 | 86.7 | 86.7 | 86.6 | 86.7 | 84.5 | 85.2 | 87.0 | 86.6 | 87.0 | 86.9 | 86.9 | 93.6 | 93.7 | 93.4 | 93.4 | 90.3 | 90.3 | 95.7 |
